# Significance filtering induces denominator selection bias in local genetic correlation: closed-form characterisation, a gate-aware correction, and an applicability diagnostic

**DOI:** 10.64898/2026.09.28.755121

**Authors:** Dana Paquin, Riddhiman Jain

## Abstract

Local genetic correlation is estimated as a ratio of a local genetic covariance to the geometric mean of two local heritabilities. LAVA, the standard implementation, returns that ratio only where both traits clear a univariate heritability test at *P <* 0.05, a rule its documentation presents as a stabiliser. We show that it is a selection event acting on the denominator, and characterise what it does to the estimates that survive. Under the noncentral Wishart model LAVA’s own interval routine assumes, the gate-conditional mean of the estimator is affine in the true correlation, with slope and intercept available in closed form from quantities every run already reports, that is, the components retained, the two univariate *P*-values, and the sample-overlap correlation. Attenuation is therefore a property of the analysis, not of the method. Across six psychiatric trait pairs the measured slope runs from 0.579 to 0.923 and the closed form predicts each to within 0.007 with nothing fitted to the panel, while within a single pair it runs from 0.371 to 0.971, so no single factor can correct it. Overlap and the filter jointly manufacture correlation where the truth is zero, most where power is lowest. We invert the closed form into a gate-aware correction, which reduces mean absolute bias from 0.247 to 0.026 over 48 simulation cells, and pair it with a diagnostic that decides whether the correction is identified. Three pairs fail it, all three sharing one trait whose local heritabilities are indistinguishable from a complete local null.

## 1 Introduction

Genome-wide genetic correlation compresses the shared genetic basis of two traits into one number, averaging over regions that may not agree. Local methods were built to recover that structure, and LAVA [1] has become the standard tool which partitions the genome into approximately independent linkage-disequilibrium blocks and estimates a genetic correlation within each. Local estimates are now read as biological findings in their own right, particularly in psychiatric genetics [2], where the question asked of them is usually about the sign of the relationship in a named region.

The estimand is a ratio that is prone to error. Its denominator is the geometric mean of the two local heritabilities, each small, noisily estimated within a single block, and free to go negative. LAVA handles this by reporting the correlation only where both traits clear a univariate heritability test at *P <* 0.05. LAVA’s documentation describes this as a way to filter out non-associated loci that may yield unstable correlation estimates.

Specifically, what it does is not benign. The univariate statistic and the denominator are built from the same sum of squares, so conditioning on the statistic being large conditions on the denominator being large. Among loci sharing a true local heritability, those admitted are disproportionately those at which sampling noise inflated the estimate, and an inflated denominator shrinks a ratio. This selection may look comparable to the winner’s curse, but the two differ in several ways. The winner’s curse selects on the estimand and inflates it [3, 4]. Here selection acts on the denominator, so the bias attenuates rather than inflates. It is affine in the true correlation rather than a threshold-dependent shift, and it interacts with sample overlap to produce non-zero expected correlation even where the truth is zero, which the winner’s curse does not.

Benchmarking work has compared local genetic correlation methods for accuracy and power under a common truth [5]. The selection induced by a method’s own reporting rule has not been characterised. We do that here, and supply a correction that inverts the characterisation with a validation of it against LAVA’s own code path on real loci, and a diagnostic that states exactly where the correction works. We then run the protocol across six trait pairs spanning the two dimensions along which the bias varies, that is, the weaker trait’s power and the overlap between the two GWAS. That survey is about how the method behaves in the regimes practitioners work in, but does not indicate or conclude anything concrete about the biology of any disorder pair.

## 2 Methods

### 2.1 The estimator, and the gate as implemented

We work from the actual LAVA source, version 0.1.5, not from its description, and state the file and line for each anomaly or inconsistency we find. Table 1 fixes notation.

**Table 1:** Notation.

| Symbol | Meaning | Source |
| --- | --- | --- |
| $K$ | principal components retained at the locus | `process.locus()`\$K |
| $\delta$ | $K \times 2$ matrix of joint effects in PC space | internal |
| $\mathbf{S} = \delta'\delta$ | sums of squares and cross-products | internal |
| $\Sigma$ | residual covariance; off-diagonal carries overlap | internal |
| $\Omega, \hat{\Omega}$ | true and estimated local genetic covariance | `locus$omega` |
| $\rho, \hat{\rho}$ | true and reported local genetic correlation | `run.bivar()`\$rho |
| $r_e$ | sample-overlap correlation between the two GWAS | cross-trait LDSC |
| $\theta_i = \Omega_{ii}/\Sigma_{ii}$ | local signal-to-noise ratio | recovered from $P_i$ |
| $U_i$ | univariate statistic, referred to $\chi_K^2$ | implied by `run.univ()`\$p |
| $G$ | the gate: both traits clear $P < 0.05$ | `analysis_functions.R:42--44` |
| $\Delta_i$ | gate-induced inflation of $S_{ii}$ | closed form, §2.3 |
| $d_i = \theta_i + \Delta_i/K$ | gated local heritability, on the $\theta$ scale | observed |
| $g_K$ | map from $\theta$ to its gated expectation | inverted in Stage 1 |
| $S_{\text{loc}}, I_{\text{loc}}$ | per-locus attenuation slope and manufactured intercept | closed form |
| $N_{\text{ref}}$ | LD reference sample size (99,339) | LAVA v1.1 panel |
| $c_{\text{ref}}$ | reference-size correction; equals 1 in v0.1.5 | `locus$nref.scale` |

For a locus retaining *K* principal components of the local LD matrix and two traits, we let ***δ*** be the *K ×* 2 matrix of joint effect estimates in principal-component space, with rows drawn independently as ***d***_*k*_ ~ *N* (***μ***_*k*_, **Σ**), the off-diagonal of **Σ** carrying the sample overlap *r*_e_. Writing ***S*** = ***δ***^*′*^***δ***, LAVA forms the bias-corrected local genetic covariance matrix as 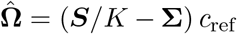 with 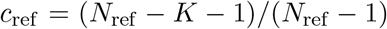 (`input_processing.R:134--166`), and the local genetic correlation as 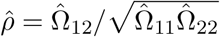 in which *c*_ref_ cancels. For binary traits the univariate statistic is *U*_*i*_ = (*S*_*ii*_*/*Σ_*ii*_) *c*_ref_, referred to 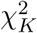. Liability-scale heritabilities are obtained afterwards [15] and play no part in the gate, which is evaluated on ***δ*** and **Σ** directly. In v0.1.5 *c*_ref_ evaluates to exactly 1, because `nref.correction` is assigned `FALSE` inside `process.locus()` and is not exposed to the caller.

Define *θ*_*i*_ = Ω_*ii*_*/*Σ_*ii*_, the local signal-to-noise ratio. Since **Σ** scales as 1*/N, θ*_*i*_ is proportional to sample size, and the univariate noncentrality is *Kθ*_*i*_, so a locus is well powered either by carrying much signal or by retaining many components. The gate is 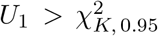 and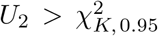 (`analysis_functions.R:42--44`). Everything below is derived for the *χ*^2^ case and so applies to binary traits only.

#### The filter and the denominator are not separable

Because 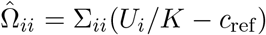 with *c*_ref_ = 1, the sign of the estimated local heritability is fixed by whether *U*_*i*_ exceeds *K*, and the 5% critical value of a 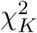 exceeds *K* for every *K*. Because passing the gate implies 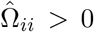 for both traits, the gate is a strict subset of the set on which 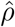 is defined at all. There is no configuration in which the filter is inactive and the estimator remains well posed, so “what LAVA would report without the filter” is not a quantity the software can produce. This determines how the attenuation must be measured, and we verify it (Section 2.5).

### 2.2 Where loci and phenotypes are lost

Tracing each loss on real data to its source identifies five distinct mechanisms, two of which are wholly undocumented and one partially (Supplementary Table S1). The consequential one is Channel 2: when a trait’s local variance estimate goes negative, `input_processing.R:169--171` silently removes that trait and returns the locus as if analysable. Analysis code assuming both phenotypes are present will therefore evaluate the gate on whichever trait survived and produce a well-formed result for a comparison that was never made, and because the trigger is a negative variance estimate the mechanism preferentially removes the weaker trait. Parsing every warning across the panel confirms the documented trigger and no second pathway: 1,407 of 1,407 drops for P3 and 1,474 of 1,474 for P5 carry a negative-variance warning. A second mechanism shows how far the filter is load-bearing: `run.bivar()` raises an error and halts on a locus it cannot analyse, and users never meet this only because the gate guarantees the positive denominator it needs.

### 2.3 The gate-conditional mean, in closed form

***S*** follows a noncentral Wishart, ***S*** *~ W*_2_(*K*, **Σ, Θ**), as confirmed by LAVA’s own interval routine, which draws from `rwishart(n.iter, K, Sigma = sigma/K, Theta = omega)`. The derivation rests on two identities that remove any need for quadrature: a noncentral *χ*^2^ is a Poisson-weighted mixture of central ones, and the truncated first moment of a central 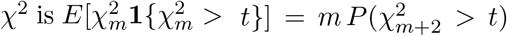 Together they give the gate pass probability and the truncated first moment of *S*_*ii*_ exactly (Supplementary Methods S1). Define Δ_*i*_ = *E*[*S*_*ii*_ | *S*_*ii*_ *> t*] *− E*[*S*_*ii*_], the gate-induced inflation of the *i*th diagonal.

Treating the numerator as unaffected gives the leading-order account, attenuation 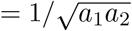 with *a*_*i*_ the conditional heritability inflation. We pre-specified that prediction and it was falsified. Simulated slopes exceed it by 8–33%. The gate inflates the numerator too, because *S*_12_ shares noise with *S*_11_ and *S*_22_. Linearising *S*_12_ on the two diagonals through the exact noncentral Wishart covariance, Cov(*S*_12_, *S*_11_) = 2*K*(*r*_e_ + Ω_12_ + *r*_e_Ω_11_) and Var(*S*_11_) = 2*K*(1 + 2Ω_11_), gives weights *b*_*i*_ = (*r*_e_ + Ω_12_ + *r*_e_*θ*_*i*_)*/*(1 + 2*θ*_*i*_) and

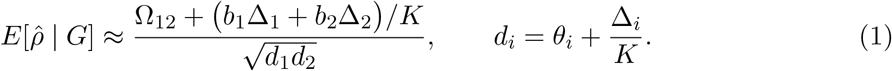

**Figure 1:**
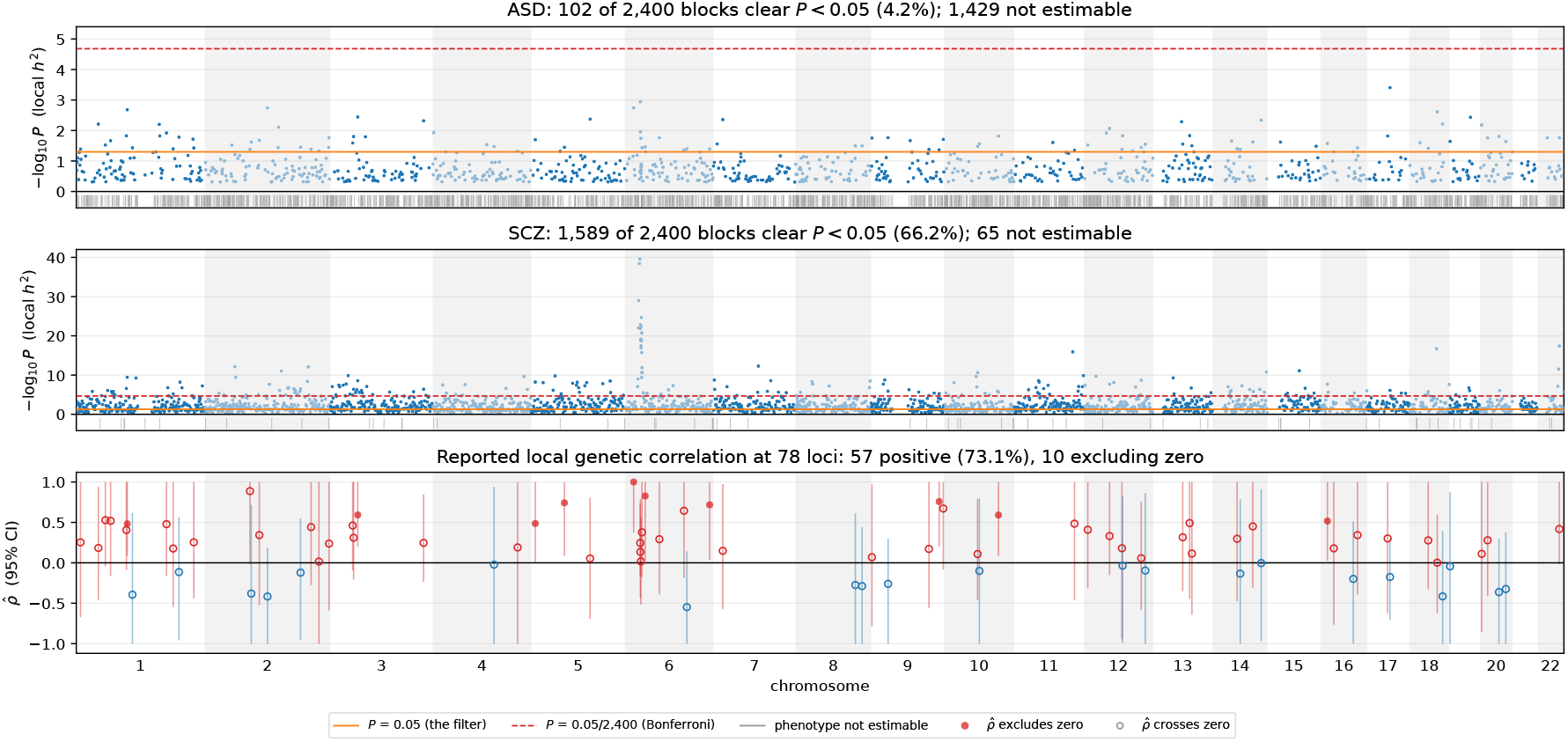
The univariate filter applied across the genome, on P4 (ASD *×* SCZ). Univariate local-heritability significance for each trait across the 2,400 processed blocks, with LAVA’s filter and the Bonferroni threshold marked, and the 78 blocks at which both traits clear the filter. Ticks below each baseline mark blocks silently dropped by Channel 2.

Each *b*_*i*_ is affine in Ω_12_, and 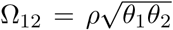 is linear in *ρ*, while Δ_*i*_ and *d*_*i*_ do not depend on *ρ* at all: the gate is a condition on the diagonals only, and their conditional moments are functions of *θ*_*i*_ and *K*. The expression is therefore affine in *ρ* exactly rather than to leading order, 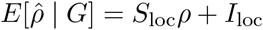 with

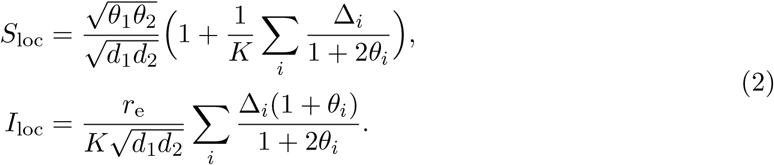

Affinity in *ρ* is algebraic rather than an expansion, that is, no term in *ρ*^2^ is dropped because none is generated, and two approximations sit behind it. The linearisation replaces *E*[*S*_12_ | *S*_11_, *S*_22_] by its least-squares linear predictor, exact for a Gaussian pair and approximate for a noncentral Wishart. And Δ_*i*_ is the inflation induced by each trait’s own gate, whereas the gate conditions on both diagonals at once: these are dependent whenever *r*_e_ ≠ 0, with 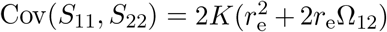, which itself carries *ρ*. That dependence vanishes at *r*_e_ = 0 and is second order in *r*_e_ elsewhere, so the residual inaccuracy is carried almost entirely by the values of *S*_loc_ and *I*_loc_ rather than by the shape. Adding a quadratic term to the pooled Tier B fit does not improve it (*P* = 0.936, Section 3.1), which bounds the departure empirically.

Two features do the work later. *d*_*i*_ is exactly the gated local heritability LAVA already reports, so the denominator 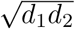 is observed, not modelled. *I*_loc_ is proportional to *r*_e_ but also to Δ_*i*_, which tends to zero as *θ*_*i*_ grows, since a locus that passes the gate almost surely is a locus on which conditioning changes nothing. The manufactured intercept is largest where power is lowest and vanishes where the filter stops binding, and *S*_loc_ *→* 1 in the same limit.

### 2.4 The correction and the applicability diagnostic

Because *S*_loc_ and *I*_loc_ are functions of quantities the analyst already holds, we invert the bias. Since 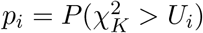 and 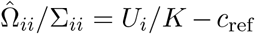 the observed gated heritability is recoverable from the reported *P*-value, and under the model 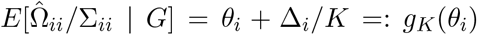 so Stage 1 de-biases the denominator. The map *g*_*K*_ is continuous and strictly increasing, so it inverts uniquely, and has a positive floor, since a locus with no true local signal still reports positive heritability once gated. That floor is the basis of the diagnostic. Stage 2 de-biases the ratio, 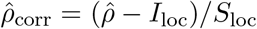. The correction inverts a conditional *mean*, and a single gated locus sits somewhere in that distribution rather than at its mean, so it must be applied to an aggregate.

#### The protocol

For the reporting loci of one trait pair: (i) read *K*, the two univariate *P*-values, 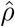 and its interval from LAVA’s output, and obtain *r*_e_ by cross-trait LD score regression; (ii) recover the observed gated heritability, *U*_*i*_ = *F* ^*−*1^(1 *− P*_*i*_; *K*) and 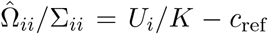 (iii) invert *g*_*K*_ on a monotone grid to recover *θ*_*i*_, stopping for any locus at or below the floor *g*_*K*_(0), where *θ* is not identified, and record it; (iv) evaluate Δ_*i*_, *d*_*i*_, *S*_loc_ and *I*_loc_; (v) pool, forming 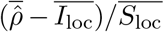 from loci-weighted means rather than inverting per locus and averaging; and (vi) transform the interval by the same affine map and report its width, which scales by 1*/S*_loc_.

#### The diagnostic

Stage 1 inverts *g*_*K*_, which flattens as *θ →* 0, so where observed gated heritability sits near the floor, a small error in the observed mean produces a large error in recovered *θ* and the correction is not identified. We report a corrected magnitude only where all three of the following checks pass, for both traits. **C1, stability**: recompute the aggregate slope under *K* stratifications into 1, 2, 3, 4, 6 and 8 strata, and pass if max */* min *<* 1.25. **C2, floor distance**: within *K* bins compare observed mean 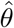 at the reporting loci against the parametric-bootstrap distribution of 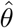 under *θ* = 0 conditional on passing the gate, and pass if pooled *z ≥* 5. **C3, the gate output is not mostly null**: from only the *P*-values compute the fraction of gate passes that are attributable to chance, referring the conditional pass rate to the conditional null 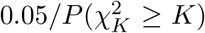 and pass if at most 0.40. C3 is included because C1 and C2 depend on the same inversion and could pass together; C3 uses only *P*-values and counting, so it cannot. Cutpoints were set on one pair (P4) and frozen before any other was computed, and every continuous quantity behind one is reported per pair. Both stages and the diagnostic are a dependency-free R module consuming exactly what LAVA already returns, so they apply to already published results without needing any re-run.

### 2.5 Simulation, and a control that attributes the effect

**Tier A** draws ***S*** directly from the noncentral Wishart and applies the gate, `param.lim` and `cap.estimates` in LAVA’s order, at 200,000 loci per cell. **Tier B** injects a known *ρ* into real loci, with their own *K*, their own **Σ** and the pair’s measured overlap, recovering it through LAVA’s actual `run.univ()` and `run.bivar()` at 2,000 replicates per cell. Tier B takes priority where the two disagree, which was decided before either was run.

**Tier C** attributes the effect, because Tiers A and B record only gated replicates and a slope below 1 measured either way is equally consistent with truncation of a heavy tail by `cap.estimates` and `param.lim`. The control is a 2 *×* 2 on a single replicate stream, so the arms are perfectly paired as *{*estimable, gated*} × {*no truncation, LAVA’s truncation*}*, where “estimable” means 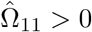 and 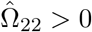. This is the weakest condition under which 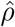 exists, and the correct no-filter baseline precisely because the gate is nested inside it. Holding truncation fixed and varying the gate isolates the filter; the reverse isolates the capping. It computes the gate statistic and 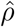 directly rather than through LAVA’s wrappers, verified against them to exact agreement over 5,000 replicates (Supplementary Methods S2). Six predictions were fixed before the runs that tested them. Three of them were confirmed and three falsified, each reported where its result appears. Correction performance was validated over 48 cells spanning *K ∈ {*100, …, 500*}*, symmetric and asymmetric heritabilities, and *r*_e_ *∈ {*0, 0.02756, 0.05, 0.20*}*, at eight true *ρ*.

### 2.6 Trait-pair panel

The protocol is applied to six pairs chosen to span the power and overlap ranges (Table 2). All of them use the UK Biobank European LD reference distributed with LAVA (v1.1, *N ≈* 99,339) and the same 2,495-block partition, so analyses are European-ancestry only, and we measure overlap per pair by cross-trait LD score regression [6, 7]. All members are case–control, which is required since the closed form is derived for the *χ*^2^ gate. P2 was designed as a controlled contrast. The PGC distributes both a full and a without-UK-Biobank release of the MDD and BIP GWAS, so P2a and P2b analyse the same two disorders with the same reference and blocks, differing only in whether UK Biobank participants are present. It did not work as designed, and we report in Section 3.2 as to why.

**Table 2:** Trait-pair panel. Pairs span the two dimensions along which the bias varies, not a disease area.

| # | Pair | Regime probed | Sources |
| --- | --- | --- | --- |
| P1 | SCZ × BIP | both well powered; moderate overlap | [9, 10] |
| P2a | MDD × BIP, full | both include UK Biobank | [11, 10] |
| P2b | MDD × BIP, no UKBB | same disorders, UKBB removed | [11, 10] |
| P3 | ASD × ADHD | high overlap, shared iPSYCH cohort | [8, 12] |
| P4 | ASD × SCZ | extreme asymmetric power; low overlap | [8, 9] |
| P5 | AN × ASD | both weakly powered | [13, 8] |

Each pair is run in its own genome pass at its own two-trait SNP intersection, because `process.input()` reduces every phenotype to the SNPs common to all phenotypes in the object. A combined pass would therefore alter the attrition cascade it exists to measure (Supplementary Methods S3). P2a and P2b are the exception as they were run together so that both arms are forced onto an identical SNP set.

## 3 Results

### 3.1 The bias is affine in *ρ*, the closed form is accurate, and the filter is its cause, not output capping

Tier B on P4 gives a pooled attenuation slope of 0.6236 (SE 0.0068, *R*^2^ 0.9527, 414 cells) with an intercept of 0.0182 (SE 0.0035, *P* = 4.3 *×* 10^*−*7^). Roughly 38% of the signal is destroyed, that is, a true local correlation of 0.50 is reported as 0.33, of which 0.31 is what the slope leaves of the signal and 0.018 the manufactured intercept. The closed form reproduces Tier A slopes to a mean relative error of 1.14% and intercepts to within 0.0052, and achieves a mean absolute error of 0.0066 on 42 held-out configurations against 0.0492 for the leading-order, denominator-only form (Supplementary Table S2). It varies with *K* at fixed heritability, from 0.4555 at *K* = 100 to 0.6897 at *K* = 500, because the noncentrality *Kθ* grows faster than the gate threshold tightens. Small blocks, which are the most numerous in the partition, have the worst attenuation, in the direction opposite to what we pre-specified. Affinity is confirmed directly by adding a quadratic term to the pooled Tier B fit, which does not improve it (*P* = 0.936).

A slope measured on gated replicates does not by itself identify what caused the attenuation, as the Tier C control does (Table 3). With `param.lim` and `cap.estimates` disabled entirely, the gated slope is still 0.647 on P4 and 0.601–0.957 across the panel, so the attenuation is fully present when nothing is truncated, and adding LAVA’s truncation to the gated arm then moves the slope by only *−*0.020 to *−*0.034 on every pair. Where the attenuation is large it is almost entirely the filter. P5, P3 and P4 lose 41.9%, 38.5% and 37.4% of the signal, of which truncation contributes 4.7%, 5.3% and 5.6%, and where truncation is a comparable contributor, on the three well-powered pairs, there is little left to attribute. Arm A is context rather than a counterfactual: its slope exceeds 1 on every pair, but on the estimable set the denominator can sit arbitrarily close to zero and 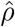 is heavy-tailed, so its mean is dominated by a few extreme replicates. The defensible statement is that among the loci LAVA reports, the estimate is attenuated, and the attenuation goes to the reporting rule.

**Table 3:**
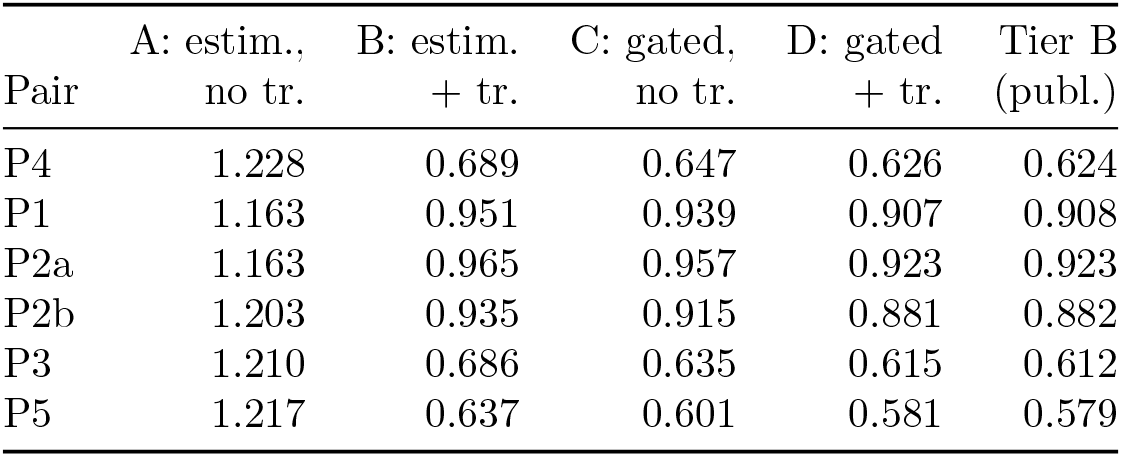
Attribution control. Slope of 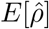 on *ρ* by selection regime, on one shared replicate stream. Arm D is what LAVA reports; “trunc.” is `param.lim` with `cap.estimates`.

The measured slope is a lower bound rather than a point estimate: 73 of 119 candidate loci were lost to Channel 2 before Tier B could use them, and each locus’s own slope regresses on the log noncentrality of its weaker trait at +0.178 (*P* = 7.7 *×* 10^*−*16^), so any Tier B estimate is fitted on a high-power subset. Per-locus slopes span 0.371 to 0.971.

**Figure 2:**
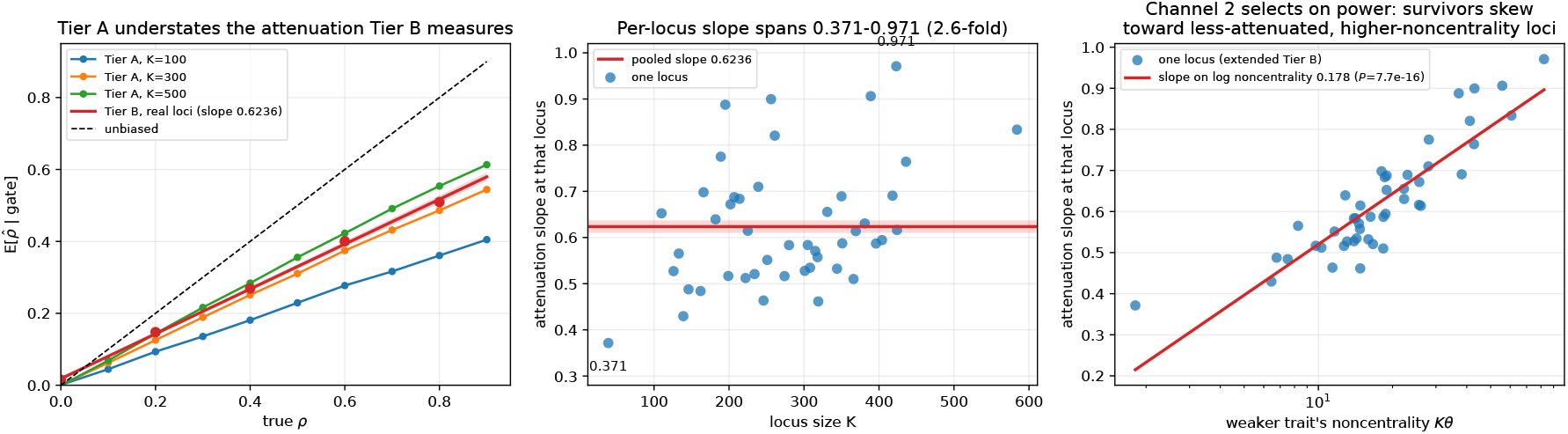
The filter attenuates the estimates that survive it. Reported 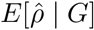 against true *ρ* under Tier A and Tier B, the fitted per-locus slope against locus size *K*, and that slope against the weaker trait’s noncentrality *Kθ*.

### 3.2 Overlap manufactures correlation, and a constant rule for it fails

At a true *ρ* of zero the filter combined with sample overlap yields a non-zero expected reported correlation. In Tier A, holding *K* = 300 and both heritabilities fixed and varying only the overlap, the reported 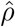 at *ρ* = 0 rises from 0.0008 at *r*_e_ = 0 to 0.0340 at *r*_e_ = 0.05 and 0.0666 at *r*_e_ = 0.10, that is, linearly in *r*_e_ with intercept *≈* 0.68 *r*_e_; a held-out cell at *r*_e_ = 0.20, with its own *K* and heritabilities, reports 0.156. On P4 that rule works: *r*_e_ = 0.02756 predicts 0.019 and Tier B on 46 real loci observed 0.0182 (*P* = 4.3 *×* 10^*−*7^), a statistically significant correlation manufactured entirely out of shared noise.

We pre-specified 0.68 *r*_e_ as the rule to be scored across the panel, but it failed by a wide margin. It predicts 0.145 on P1 where Tier B measures 0.028, and averaged over six pairs its absolute error is 0.0432, which on a parameter bounded in [*−*1, 1] is not a usable prediction. The constant was read off Tier A in a low-power configuration, but *I*_loc_ *∝* Δ_*i*_ and Δ_*i*_ *→* 0 as power grows, so the manufactured intercept must shrink for well-powered pairs, which are exactly the pairs with the largest *r*_e_ in this panel. Evaluating the closed form on each pair’s own per-locus (*K, θ*_1_, *θ*_2_, *r*_e_) reduces the mean absolute error from 0.0432 to 0.0050 (Table 6).

The replacement statement is conditional. Where the filter binds, overlap manufactures correlation at a magnitude that can exceed published local genetic correlations: 0.139 on P3, whose ASD and ADHD GWAS share the iPSYCH cohort. Where it is close to non-binding the intercept is small almost regardless of *r*_e_: P1 has the panel’s largest overlap at *r*_e_ = 0.214 and one of its smallest intercepts at 0.028. Overlap alone is insufficient for an analyst to know whether there is a problem, but together with power it is informative enough. The P2 contrast exposed this as removing UK Biobank did not lower the measured overlap (0.1090 *→* 0.1132) but did lower power, and the closed form tracks the resulting rise in both arms’ intercepts because it is a function of both jointly.

### 3.3 Intervals are calibrated and nearly vacuous

Post-filter coverage is 0.949 against a nominal 0.950, so the filter does not break LAVA’s intervals. Width is however a problem. Mean simulated post-filter width is 1.386 on a parameter bounded in [*−*1, 1], spanning 69% of the parameter space, and the simulation predicts the real intervals closely without being fitted to them (1.365 observed on P4). The fraction of simulated intervals excluding zero is 0.139 at *ρ* = *−*0.4 and 0.123 at *ρ* = +0.4, so the count of sign-determinate loci cannot discriminate between hypotheses about direction. On P4 every one of the ten determinate intervals has its upper bound pinned at 1.000 by `cap.estimates`, so the sign is determined while the magnitude is not.

**Figure 3:**
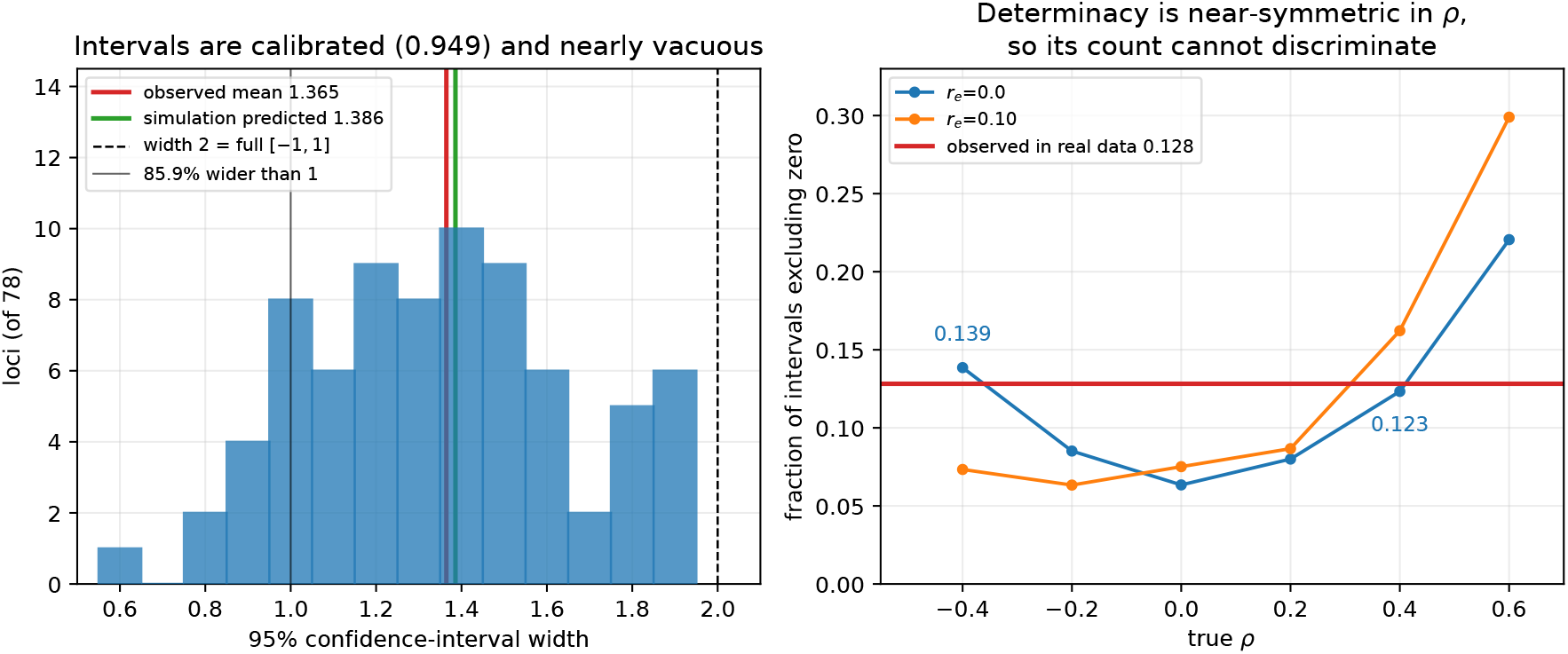
The intervals are correctly calibrated and nearly uninformative. Distribution of 95% confidence-interval width across P4’s 78 reported loci, against the width-2 line that marks the whole parameter space, and the fraction of intervals excluding zero as a function of true *ρ*.

**Figure 4:**
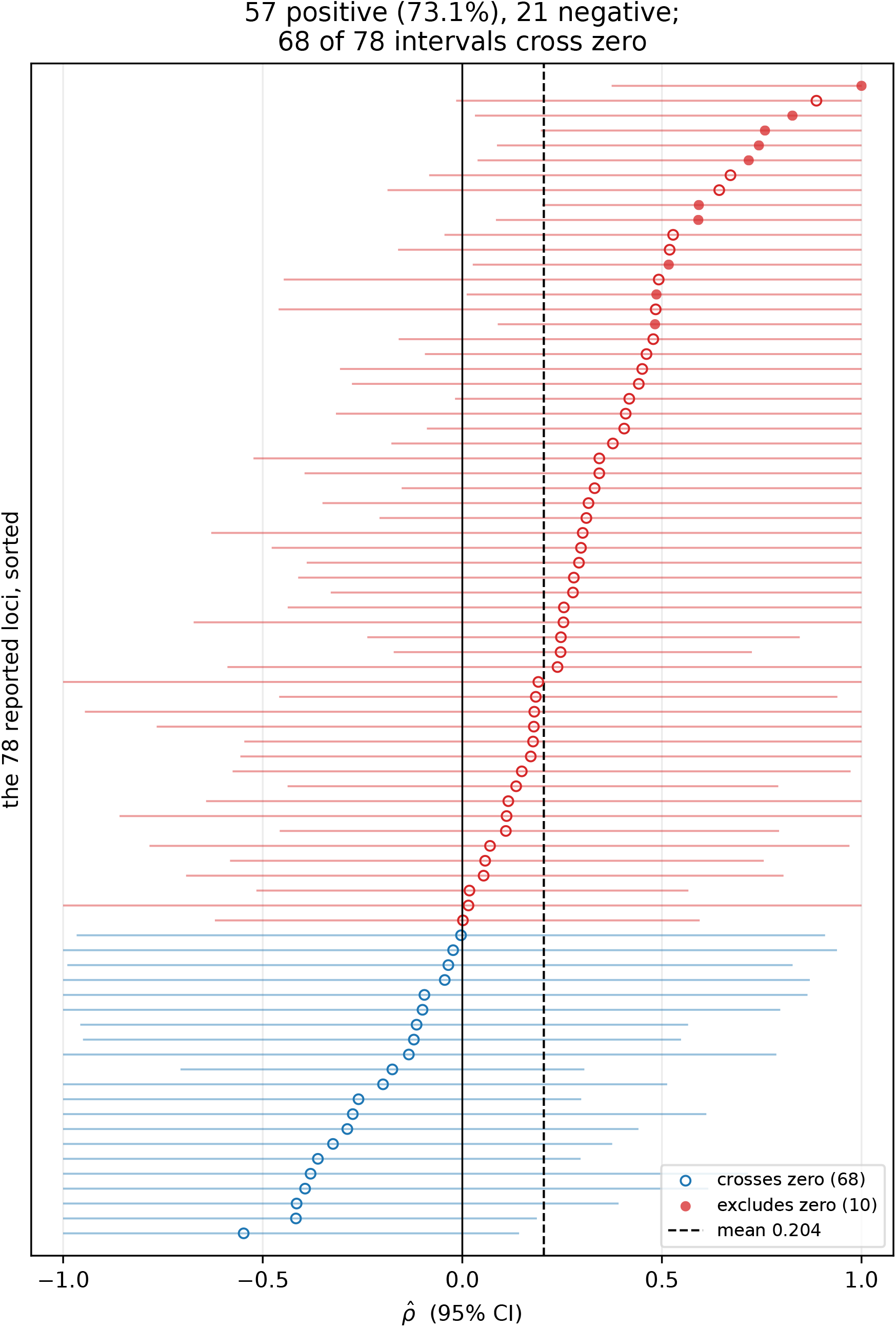
All 78 reported local genetic correlations for P4, sorted. Each row is one locus with its 95% confidence interval, coloured by sign; filled points are the 10 intervals excluding zero and open points the 68 crossing it.

### 3.4 The correction, and the penalty for applying it per locus

Over 48 simulation cells the aggregate gate-aware correction reduces mean absolute bias from 0.247 to 0.026, a factor of 9.7, and by a factor of 7.6 relative to a single pooled factor (Table 4). The pooled factor uses the best available genome-wide slope, 0.6236, and performs worse than the earlier and less accurate 0.5512 did (0.195 against 0.182). A single factor fails to encapsulate the heterogeneity it is asked to summarise. A pooled factor is not an approximation to the right answer that improves as the pooling improves, it is the wrong shape of answer.

**Table 4:** Correction performance over 48 simulation cells.

| Estimator | Mean abs. bias | Max abs. bias |
| --- | --- | --- |
| Uncorrected, as LAVA reports | 0.247 | 0.602 |
| Single pooled factor ($\div 0.6236$) | 0.195 | 0.579 |
| Gate-aware, applied per locus | 0.106 | 0.304 |
| Gate-aware, applied in aggregate | 0.026 | 0.107 |

The per-locus variant is markedly worse than the aggregate one, 0.106 against 0.026, because *g*_*K*_ is nonlinear and floored. Correction also does not rescue per-locus inference: the interval transforms by the same map, so its width scales by 1*/S*_loc_, between 1.95*×* and 3.01*×* across the validation cells. Applied to the observed mean width of 1.365 that gives 2.27, wider than the entire parameter space. Correcting a point estimate cannot narrow its interval, and the interval is the binding constraint on any per-locus claim.

**Figure 5:**
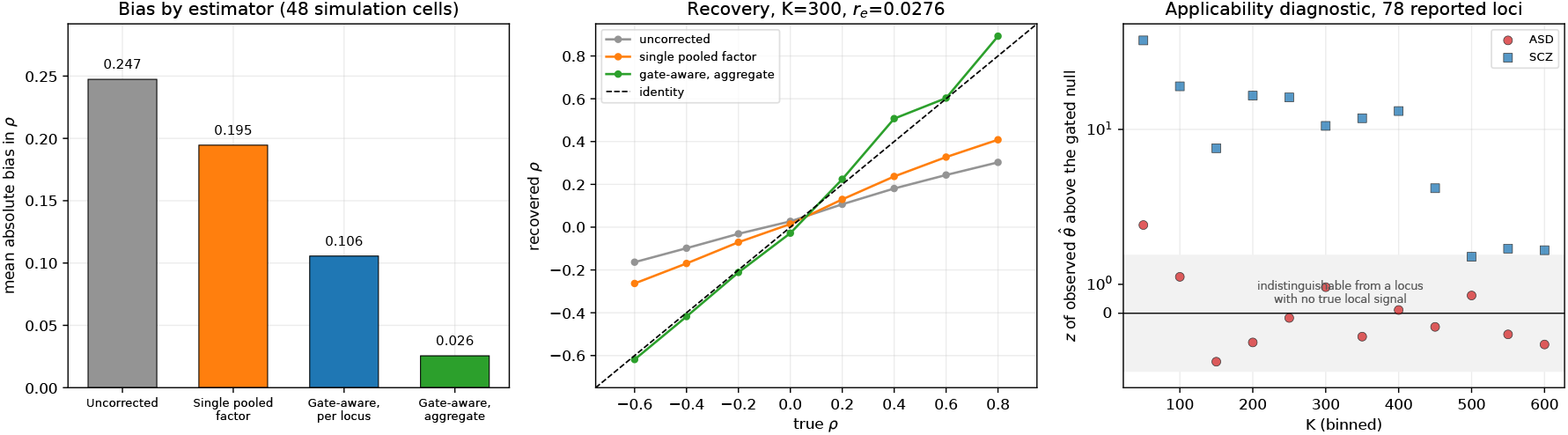
The correction and its diagnostic. Mean absolute bias and recovery of the true correlation under four estimators across the 48 validation cells, and the applicability diagnostic on real data by *K* bin and trait, with the *±*2 band marking indistinguishability from a locus carrying no true local signal.

### 3.5 Three loci through the protocol

Three loci from P1, the panel’s best-behaved pair, show the three cases the protocol has to distinguish (Table 5). At locus A the weaker trait clears the filter but not comfortably (*P* = 0.011), so the gate is doing real selection work and the slope is 0.7217. The corrected estimate nonetheless moves only from 0.2588 to 0.2711, because at *r*_e_ = 0.2139 the filter manufactures +0.0632 of apparent positive correlation here and removing it before dividing all but cancels the inflation from dividing. A practitioner who divided by the slope and forgot the intercept would report 0.359 instead of 0.271, a 32% overstatement, which is a likely way to misapply the correction. At locus B the correction is inert and should be, as both traits clear the filter by orders of magnitude, the slope is 0.9881, and 0.4922 becomes 0.4916.

**Table 5:** Three P1 reporting loci taken through the correction; *r*_e_ = 0.2139.

|  | A: loc. 874 | B: loc. 38 | C: loc. 168 |
| --- | --- | --- | --- |
| Chromosome, $K$ | chr5, 357 | chr1, 285 | chr1, 345 |
| $P$, weaker trait | 0.0114 | $1.78 \times 10^{-4}$ | 0.0310 |
| $P$, stronger trait | $5.1 \times 10^{-11}$ | $1.34 \times 10^{-5}$ | $1.49 \times 10^{-5}$ |
| $\hat{\rho}$ as reported | 0.2588 | 0.4922 | -0.6064 |
| Recovered $\theta_1$ | 0.0800 | 0.3192 | 0.0000 |
| Recovered $\theta_2$ | 0.5660 | 0.3913 | 0.3509 |
| $S_{\text{loc}}$ | 0.7217 | 0.9881 | — |
| $I_{\text{loc}}$ | 0.0632 | 0.0064 | — |
| At the selection floor? | No | No | Yes |
| $\hat{\rho}_{\text{corr}}$ | 0.2711 | 0.4916 | not identified |
| Interval width, after | 1.3754 | 0.8656 | — |

**Table 6:** Protocol output across the panel. Closed-form slope and intercept are *S*_loc_ and *I*_loc_ evaluated on each pair’s own per-locus (*K, θ*_1_, *θ*_2_, *r*_e_) and averaged over the loci Tier B fitted; measured values are that pair’s extended Tier B fit. C1–C3 are the diagnostic checks of Section 2.4, reported as frozen. Percentages for estimability and Channel 2 loss are of processed blocks.

| Quantity | P4 | P1 | P2a | P2b | P3 | P5 |
| --- | --- | --- | --- | --- | --- | --- |
| Blocks processed | 2,400 | 2,487 | 2,483 | 2,483 | 2,424 | 2,458 |
| Weaker trait estimable | 40.5% | 95.3% | 86.7% | 74.4% | 42.0% | 40.0% |
| Both estimable | 37.8% | 92.7% | 83.9% | 72.9% | 40.3% | 39.1% |
| Reported loci | 78 | 1,369 | 816 | 545 | 84 | 73 |
| Reported, % of blocks | 3.13 | 54.87 | 32.71 | 21.84 | 3.37 | 2.93 |
| Bonferroni survivors | 0 | 98 | 20 | 8 | 0 | 0 |
| Channel 2 loss, weaker | 59.5% | 4.7% | 13.3% | 25.6% | 58.0% | 60.0% |
| Channel 2 loss, stronger | 2.7% | 2.6% | 3.1% | 2.0% | 1.7% | 0.9% |
| Median $K$ | 268 | 295 | 290 | 290 | 229 | 223 |
| Measured $r_e$ | 0.0276 | 0.2139 | 0.1090 | 0.1132 | 0.2372 | 0.0945 |
| Slope, closed form | 0.620 | 0.902 | 0.917 | 0.877 | 0.619 | 0.578 |
| Slope, measured (Tier B) | 0.624 | 0.908 | 0.923 | 0.882 | 0.612 | 0.579 |
| Intercept, closed form | 0.0174 | 0.0360 | 0.0166 | 0.0249 | 0.1476 | 0.0696 |
| Intercept, measured | 0.0182 | 0.0283 | 0.0125 | 0.0205 | 0.1389 | 0.0652 |
| Intercept, 0.68 $r_e$ rule | 0.0187 | 0.1454 | 0.0741 | 0.0770 | 0.1613 | 0.0643 |
| C1, slope fold-change | 2.66 | 1.02 | 1.03 | 1.08 | 2.87 | 4.70 |
| C2, pooled $z$, weaker trait | −0.2 | +38.8 | +18.0 | +8.1 | −0.2 | −0.2 |
| C3, null fraction, weaker | 0.49 | 0.07 | 0.10 | 0.14 | 0.44 | 0.53 |
| Corrected magnitude usable? | No | Yes | Yes | Yes | No | No |

Locus C cannot be corrected at all. Inverting *g*_*K*_ returns *θ*_1_ = 0 exactly, so no *θ* is consistent with the data other than the floor. This is not a rare edge case, as 353 of P1’s 1,369 reporting loci (25.8%) sit at the floor, in the pair that passes the diagnostic most comfortably in the panel. A quarter of the elements do not exist, which is the reason the correction is pooled.

### 3.6 The panel: attrition, prediction and diagnostic

All six pairs complete (Table 6). The rows carrying the most weight are the closed-form and measured slopes, the only genuinely out-of-sample test in the paper: the closed form is derived from a sampling model, while the measured value comes from injecting a known *ρ* into real loci and recovering it through LAVA’s own code. The largest discrepancy is 0.007 and the mean 0.0048, and the agreement holds across a slope range of 0.579 to 0.923, from a regime where the filter destroys 42% of the signal to one where it costs 8%, with no parameter fitted to the panel. That is what makes the protocol usable. An analyst holding *K*, two univariate *P*-values and an LDSC overlap estimate can compute the expected attenuation of their own analysis before deciding whether to trust it.

The dominant loss is not the filter. On P4 the weaker trait’s local variance estimate goes negative at 1,429 of 2,400 processed loci (59.5%) against 65 (2.7%) for the stronger, due to the undocumented Channel 2. While we had expected the filter to be the primary constraint, the weaker trait sets the ceiling at every stage.

The diagnostic separates the panel cleanly. P5 reports 73 loci and P4 reports 78, while accepted P2b reports 545, so the separation is not by effect size or locus count. It detects whether the denominator carries signal at all. One correction to our own procedure belongs here: C3 as frozen referred the conditional pass rate to a flat *α* = 0.05 rather than to the conditional null 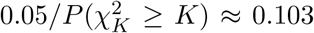 so every value roughly doubles when recomputed. The 0.40 threshold falls between the two groups under either baseline and all six verdicts are identical. We report both.

### 3.7 A pair that fails the diagnostic, and what that shows

P4 fails all three criteria, and the failure is instructive because it is the regime a large fraction of psychiatric applications occupy. Across *K* stratifications from 1 to 8 strata its aggregate slope moves from 0.769 to 0.288 and the implied corrected mean from 0.25 to 0.64. The floor diagnostic explains why. At the 78 reporting loci the weaker trait’s observed mean gated heritability is indistinguishable from what a locus with no true local signal reports after passing the filter, at per-bin *z* from *−*1.67 to +3.05 against +1.95 to +48.3 for the stronger trait.

**Figure 6:**
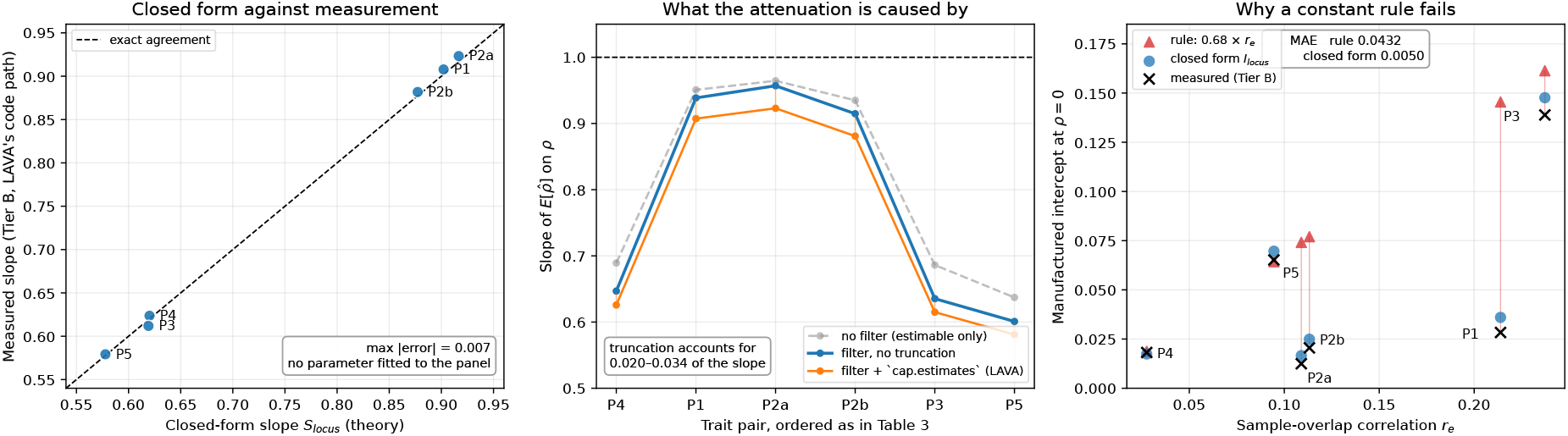
Validation and attribution. (a) The closed-form slope *S*_loc_ against the slope Tier B measures on each pair’s real loci, for all six pairs; (b) the attribution control of Table 3; (c) the manufactured intercept against *r*_e_, with the pre-specified 0.68 *r*_e_ rule, the closed form and the measurement.

A test that does not depend on the pass rate at all agrees, which is decisive. Under a complete local null *p*_*i*_ is uniform and Channel 2 retains exactly those loci with 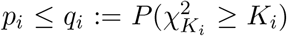, so *u*_*i*_ = *p*_*i*_*/q*_*i*_ is Uniform(0, 1) on the retained loci. Across the weaker trait’s 971 estimable loci that null is not rejected: mean *u* = 0.491 against 0.5, Kolmogorov–Smirnov *P* = 0.47. The same trait gives *P* = 0.078 and *P* = 0.86 in the two other pairs it appears in, while all nine remaining trait-analyses in the panel reject at *P <* 10^*−*120^.

We conclude that within this analysis the weaker trait carries no detectable local heritability signal anywhere in the genome, and the 78 reported loci are not a set of weak findings but what the filter emits when one of its two inputs is noise. Taking them at face value compounds the error. 57 of 78 are positive, which a binomial test against 50/50 returns at *P* = 5.6*×*10^*−*5^, but 0.5 is not the null when the two GWAS share samples, and the filtered null gives *P* = 3.8 *×* 10^*−*4^. Dividing such estimates by a genome-wide attenuation slope, the only correction previously available, actively misleads, because the reporting loci are the high-power tail where attenuation is mildest.

## 4 Discussion

### 4.1 Reporting guidance

We suggest some guidance on reporting that users of LAVA, and of comparable tools, can follow, each item traceable to a result.

1. **Report the attrition cascade, not only the reported loci**: how many blocks entered, how many were processed, how many were estimable per trait, and how many passed. A study reporting 78 loci without stating that the other 2,417 blocks returned no estimate misleads the reader into assuming that the genome was tested.
2. **Check for Channel 2 explicitly**. Verify that `process.locus()` returned both pheno-types before scoring the filter. Code that does not check this will report a comparison that was never made.
3. **Never correct with a single factor**. The slope varies from 0.371 to 0.971 across real loci within one pair and from 0.579 to 0.923 between pairs. Compute *S*_loc_ instead.
4. **Run the applicability diagnostic before reporting any corrected magnitude**, and where observed gated heritability is not clear of the selection floor, report the uncorrected estimate with the attenuation stated as a bound.
5. **Report interval width alongside coverage, and do not test sign against 50/50**. Coverage is fine, but width is the problem. Where the two GWAS share samples, the filtered null is shifted, so the naive test overstates significance.
6. **Compare conditional rates against conditional nulls**. A gate pass rate computed among loci that survived Channel 2 must be referred to 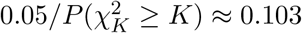 not to 0.05. We made this error ourselves.
7. **Report the phenotype set used to build the input object, and state** *r*_e_ **without reading the manufactured intercept off it alone**. That composition sets the *K* cap and the common-SNP set, and the panel’s highest-overlap pair (*r*_e_ = 0.214) has one of its smallest intercepts (0.028).

### 4.2 What this means for already published results, and how far it generalises

The correction consumes only *K*, two univariate *P*-values, 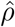 with its interval, and an overlap estimate, each printed in a typical LAVA supplementary table or recoverable from one, so the protocol applies retrospectively without the original genotypes, the LD reference, or the summary statistics. That was a design requirement, because the great majority of local genetic correlations that will ever be interpreted have already been published.

Most published estimates are probably fine, and this paper should not be read as saying otherwise. Attenuation tracks the weaker trait’s power and on the panel ranges from 8% to 42%. A study of two well-powered disorders with large *K* is in P1’s regime, where the slope is 0.908 and the manufactured intercept 0.028 despite an *r*_e_ of 0.214. Its bias is a function of the analysis, is currently unreported, and is now computable. Where the weaker trait is underpowered the magnitudes should be treated as lower bounds and the sign claims re-examined, and that regime is recognisable without re-analysis: a large Channel 2 loss rate for one trait, a small reporting fraction, and few or no Bonferroni survivors.

The closest classical relative is Berkson’s admission artefact [14], in which a sample selected on a function of two variables acquires an association absent in the population. The difference is that the selected variables here are the estimand’s denominator, so the induced dependence propagates into a ratio. That structure is not specific to LAVA, and whether *ρ*-HESS, SUPERGNOVA and LOGODetect share it is a question the benchmarking literature [5] is well placed to answer and did not ask.

### 4.3 Limitations

- **The attenuation slope is a lower bound, not a point estimate**. Tier B can be fitted only on loci surviving Channel 2, which removes low-power loci, and enlarging the sample does not fix it, so no unbiased genome-wide slope is obtainable this way.
- **A two-trait local genetic correlation is not reproducible from the trait pair alone**. `process.input()` reduces every phenotype to the SNPs common to all phenotypes in the object: analysing ASD *×* SCZ inside an eight-phenotype object would have cut usable SNPs from 5,774,835 to 2,306,339. Since the slope depends strongly on *K*, the same two GWAS analysed in a larger object return a more attenuated answer, and published estimates from multi-phenotype studies are not directly comparable to two-trait ones.
- **The** 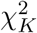 **reference distribution is not exactly calibrated on these data, and we cannot fully explain the discrepancy**. A complete local null implies survival of Channel 2 at 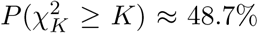 of loci and real signal can only raise that figure, yet one trait sits below it in all three analyses it appears in (40.5%, 42.0%, 40.0%). Verdicts are unaffected, but the absolute calibration of the floor quantities is approximate.
- The derivation covers binary traits, since the *F*-test LAVA applies to continuous traits changes the selection geometry, and analyses are European-ancestry only, forced by the LD reference.
- The second-order form is an approximation, though the discrepancy is at most 0.007 against Tier B, and the diagnostic is a necessary condition rather than a sufficient one: passing it only establishes that *θ* is identified, not that the sampling model holds.

## 5 Conclusion

LAVA’s univariate significance filter is a selection event on the denominator of a ratio estimator, and the two are not separable: passing the filter implies a positive denominator, and `run.bivar()` cannot run without one. The filter’s effect is available in closed form under LAVA’s own sampling model, affine in the true correlation, and computable per locus from quantities every run already reports. Across six trait pairs that closed form predicts the measured attenuation slope to within 0.007, with nothing fitted to the panel.

The size of the bias is a property of the analysis. Where the filter binds the damage compounds: 42% of the signal destroyed, per-locus slopes from 0.371 to 0.971 so that no single factor can correct it, an overlap-driven intercept manufacturing significant correlation from shared noise, and intervals too wide to determine the sign at 87% of the loci it reports (68 of 78 on P4). Where it does not bind, LAVA is close to unbiased and the correction here close to unnecessary. That is itself worth reporting, because the alternative reading of an uncharacterised filter is that every published estimate is suspect, and that is not what we find. A study that reports filtered local genetic correlations without the attrition cascade, the interval widths and, where sign is at issue, the filtered null is not reporting a genome-wide test, but the small, power-selected window its weaker trait permitted.

## Acknowledgments

The authors thank the Psychiatric Genomics Consortium and the contributing study participants, whose summary statistics made this work possible. The authors also thank Dr. Alex Becker for mentorship and guidance throughout this project, gratefully acknowledged for his support throughout the project.

## Use of AI tools

The author used large language models to edit the text of this manuscript and to help write the analysis code. No section of the text, and no figure, table or reference, was taken from a model’s output without the author’s own revision. All code was read and checked by the author, and every numeric claim in the manuscript is verified against its source file by `code/python/16 audit numbers.py`. The author is responsible for the whole of the content.

## Data availability

All analysis code, the correction module and every result file underlying a number in this paper are deposited at https://github.com/riddhimanjain/lava-gate and archived at Zenodo under the MIT licence, at https://doi.org/10.5281/zenodo.22830134 for the version used here and https://doi.org/10.5281/zenodo.22132758 for all versions. The deposit is self-contained: the verification and audit scripts of Sections 2.5, 3.1, 3.2 and 3.7 run from the deposited `results/` directory alone, without the bulk data.

The correction and the applicability diagnostic are supplied as a dependency-free R module (`code/R/lavagate.R`, base R *≥* 4.0) consuming LAVA’s own output, so they can be applied to previously published local genetic correlations without re-running any analysis. Two tests are included and are intended to be run before the code is trusted. `code/R/test_lavagate.R` checks the R implementation against the Python one used for this paper’s numbers; they agree to 1.3*×*10^*−*12^ on Δ and 1.8*×*10^*−*15^ on the slope and intercept over 500 parameter combinations spanning *K ∈* [50, 500], *θ ∈* [0.005, 0.30] and *r*_e_ *∈* [0, 0.20], and it re-derives P4’s published aggregate slope, intercept and interval widths. `code/python/14_pair_pipeline.py --selftest` checks that the generic per-pair pipeline reproduces P4 to floating point, including its diagnostic verdict.

Verification is deposited as runnable code and as raw output under `results/verification/`: `V0_exactness_check.R` (agreement with `run.univ()` and `run.bivar()`, and the nesting of Section 2.1), `V1_gate_attribution.R` (Table 3), `V2_gate_null baseline.py` and `V3_null_calibration.py` (the conditional null and the complete-null goodness of fit), `V4_closedform_vs_tierB.py` (Table 6) and `V5_null_recalibration.py` (the calibration limitation of Section 4.3). V0 is a precondition for V1 and exits non-zero if agreement is not exact. Every numeric claim in this manuscript is checked against its source file by `code/python/16_audit_numbers.py`.

GWAS summary statistics are publicly available from the Psychiatric Genomics Consortium (https://pgc.unc.edu/for-researchers/download-results/) and are documented with accession details and download locations in `data/DATA_SOURCES_MULTIPAIR.md` rather than redeposited. The LAVA LD reference panel (v1.1) and the 2,495-block genome partition are distributed with LAVA. Reproducing the panel end to end additionally requires those bulk files (approximately 16 GB). Software versions: LAVA 0.1.5, R 4.3.1, Python 3.10.12, `numpy` 1.24.4, `scipy` 1.10.1.

## Supplementary

Supplementary material is provided as a separate file, and contains Supplementary Methods S1–S3, Supplementary Tables S1–S3 and Supplementary Figure S1.

