## Supplemental Data 1 for "Significance filtering induces denominator selection bias in local genetic correlation: closed-form characterisation, a gate-aware correction, and an applicability diagnostic"

This document contains Supplementary Methods S1–S3, Supplementary Tables S1–S3 and Supplementary Figure S1, referenced from the main text. Every number is generated by the deposited code; none is entered by hand.

**Use of AI tools.** The authors used large language models to edit the text of this document and to help write the verification code referenced throughout (for example `code/replication/V0_exactness_check.R` and `code/R/test_lavagate.R`). No section of the text, and no table or figure, was taken from a model's output without the authors' own revision. All code was read and checked by the authors, and every number reported here is regenerated by the deposited scripts rather than entered by hand. The authors are responsible for the whole of the content.

#### Contents

**S1 The exact moment identities, and their implementation** 1

**S2 Verification of the attribution control** 2

**S3 One genome pass per pair, and why not one pass for the panel** 2

**S4 Supplementary tables** 3

**S5 Supplementary figure** 5

#### S1 The exact moment identities, and their implementation

The derivation of Section 2.3 of the main text needs two quantities for a noncentral $\chi^2$ variate: the probability that it exceeds the gate threshold, and its first moment truncated to that region. Both are available in closed form, so the correction contains no numerical integration.

**The Poisson mixture.** A noncentral chi-square with $m$ degrees of freedom and noncentrality $\lambda$ is a Poisson-weighted mixture of central chi-squares,

$$\chi'^2_m(\lambda) \;=\; \sum_{j\geq 0} \mathrm{Pois}(j;\lambda/2)\,\chi^2_{m+2j}, \tag{S1}$$

so any functional of the noncentral law that is available for the central law is available for the noncentral one by summing a rapidly convergent series. In our implementation the series is truncated where the weight falls below $10^{-14}$ of its maximum, with the index bounded by $\lambda/2 + 12\sqrt{\lambda/2}$.

**The truncated first moment.** For a central chi-square,

$$E\big[\chi^2_m \, \mathbf{1}\{\chi^2_m > t\}\big] \;=\; m\, P(\chi^2_{m+2} > t), \tag{S2}$$

which follows in one line by absorbing the extra factor of $x$ into the density. Since $f_m(x) \propto x^{m/2-1}e^{-x/2}$, we have $x\, f_m(x) = m\, f_{m+2}(x)$.

Combining the two gives the gate pass probability and the truncated first moment of $S_{ii}$ exactly. Writing $t = \chi^2_{K,\,0.95}$ and $\lambda_i = K\theta_i$,

$$\Delta_i \;=\; E[S_{ii} \mid S_{ii} > t] - E[S_{ii}] \tag{S3}$$

is the gate-induced inflation of the $i$th diagonal used throughout the main text. It conditions on trait $i$'s own gate. The reported gate conditions on both, and the two diagonals are dependent whenever $r_\mathrm{e} \neq 0$, with $\mathrm{Cov}(S_{11}, S_{22}) = 2K(r_\mathrm{e}^2 + 2r_\mathrm{e}\Omega_{12})$, so $\Delta_i$ is exact at $r_\mathrm{e} = 0$ and accurate to second order in $r_\mathrm{e}$ otherwise.

**The reference-size correction.** $c_\mathrm{ref} = (N_\mathrm{ref} - K - 1)/(N_\mathrm{ref} - 1)$ is written, named and applied at three separate points of the LAVA source, but `nref.correction` is assigned `FALSE` inside the body of `process.locus()` (`input_processing.R:38`), is not exposed as a parameter, and cannot be set by the caller. In v0.1.5 it therefore evaluates to exactly 1 and the correction is inert. We retain the general form in the main text because it is what the code expresses and what a reimplementation would apply. At $K = 270$ and $N_\mathrm{ref} = 99{,}339$ the intended factor is 0.99727, so the distinction shifts the gate threshold by 0.27% and no quantity reported in the paper depends on it. Tier B, running through LAVA's own code path, already reflects $c_\mathrm{ref} = 1$.

**Cross-implementation check.** The R module `code/R/lavagate.R` and the Python used for every number in the paper are independent ports of the same closed form. They are checked against each other on 500 parameter combinations spanning $K \in [50, 500]$, $\theta \in [0.005, 0.30]$ and $r_\mathrm{e} \in [0, 0.20]$, agreeing to $1.3 \times 10^{-12}$ on $\Delta$ and $1.8 \times 10^{-15}$ on the slope and intercept (`code/R/test_lavagate.R`).

#### S2 Verification of the attribution control

The Tier C control of Section 2.5 evaluates the gate statistic and $\hat\rho$ directly rather than through `run.univ()` and `run.bivar()`, which would be infeasible at the replicate count required. That substitution is legitimate only if the direct formulas are LAVA's, so we verify it rather than assert it, by pushing identical $\boldsymbol{\delta}$ matrices through both paths and comparing them.

Over 5,000 replicates spanning $K \in [39, 584]$ and $r_\mathrm{e} \in [0, 0.24]$, agreement is exact, at a maximum relative difference of 0 on all 10,000 univariate $P$-values and all 1,909 defined $\hat\rho$ values, after applying the `signif(., 6)` rounding LAVA applies to its own output. The same run confirms the nesting of main-text Section 2.1 empirically. Of 148 gate passes, zero had a non-positive denominator, and all 2,322 `run.bivar()` aborts were loci that were not estimable.

Because the simulation depends on a locus only through $(K, \theta_1, \theta_2, r_\mathrm{e})$, since both $\hat\rho$ and $U_i$ are invariant to rescaling $\boldsymbol{\Sigma}$ and $\boldsymbol{\Omega}$ together, the control uses the real per-locus values measured by Tier B, at twice its replicate count. The verification is deposited as `code/replication/V0_exactness_check.R`, which is a precondition for `V1_gate_attribution.R` and exits non-zero if agreement is not exact.

#### S3 One genome pass per pair, and why not one pass for the panel

`process.locus()` returns $\boldsymbol{\Omega}$ and $\boldsymbol{\Sigma}$ for every phenotype in its input object, so a single genome pass over all eight phenotypes would in principle serve all six pairs at a sixth of the compute.

Three mechanisms make $K$ depend on the phenotype set, and the third is decisive.

Two do not bind at this panel's sample sizes. The cap `max.K` $= \lfloor$`max.prop.K` $\cdot \min(N)\rfloor$ evaluates to 34,762 for the smallest study (ASD, $N$ = 46,350) against a largest observed $K$ of 607, and the rule that nulls $h^2$ when $K/N > 0.1$ is far from triggered at $K/N = 0.013$.

The third binds. `process.input()` reduces every phenotype to the SNPs common to *all* phenotypes in the object and the reference panel. Measured on the harmonised panel, ASD $\times$ SCZ with the reference retains 5,774,835 SNPs, whereas all eight phenotypes retain 2,306,339. Since $K$ is close to proportional to the number of SNPs in a block, and the univariate noncentrality is $K\theta$, a combined pass would not merely be less precise. It would alter the attrition cascade it exists to measure, and the completed pair would not reproduce its own published numbers.

Each pair is therefore run in its own pass, at its own two-trait SNP intersection, with one exception. P2a and P2b are run together, in a single four-phenotype pass, because the two PGC releases have genuinely different SNP coverage and running the arms separately would add a second difference between them and destroy the control. Sharing one pass forces the identical SNP set on both arms, at the cost of a smaller SNP set for P2 alone. Because `process.locus()` drops phenotypes individually, a locus in that pass can be analysable for one arm and not the other, so the surviving phenotype set is recorded per locus and each arm is scored independently against it.

#### S4 Supplementary tables

Table S1: **Where loci and phenotypes are lost.** Five mechanisms, identified by tracing each loss on real data back to the LAVA v0.1.5 source. Channels 2 and 5 are wholly undocumented; channel 4 is documented in part.

| # | Mechanism | Location and effect | Documented |
|---|---|---|---|
| 1 | Locus not processed | `process.locus()` returns `NULL`: insufficient SNPs after harmonisation, or *all* phenotypes have negative local variance | Yes |
| 2 | Phenotype discarded | `input_processing.R:169--171`: negative local variance for one trait, or an `NA` from $N < K$ or inversion failure; that trait is silently removed and the locus returned as if analysable | **No** |
| 3 | The univariate filter | `analysis_functions.R:42--44`: both traits must clear $P < 0.05$ | Yes |
| 4 | Parameter truncation | `analysis_functions.R:384--385`: $\hat{\rho}$ set to `NA` when $\lvert\hat{\rho}\rvert >$ `param.lim` (default 1.25); survivors then capped to $[-1, 1]$ at line 174 | Partly |
| 5 | `run.bivar()` aborts | `analysis_functions.R:375`, then 387: on a locus with a negative local variance, `estimate.params()` returns `NaN` and `filter.params()` evaluates `if (any(NA))`, raising *"missing value where TRUE/FALSE needed"* | **No** |

Channel 2 is the consequential one and is quantified per pair in main-text Table 6. Channel 5 is reported for what it implies rather than for how often it bites. `run.bivar()` does not return `NA` for a locus it cannot analyse, it raises an error and halts. Users never meet this, because `run.univ.bivar()` only reaches the bivariate step through the filter, and the filter guarantees a positive denominator. The filter is therefore load-bearing for the pipeline to run at all, not an optional stabiliser that could simply be switched off. Channel 4 is a filter on the answer rather than on the data, and its two stages are often conflated. `param.lim` discards $\lvert\hat{\rho}\rvert > 1.25$ entirely, while `cap.estimates` silently rewrites $|\hat{\rho}|$ between 1 and 1.25 to exactly $\pm 1$.

Table S2: **Closed form against Tier A Monte Carlo**, 200,000 loci per cell. Mean relative error on the slope is 1.14% and the maximum 2.06%; maximum absolute error on the intercept is 0.0052. The closed form returns an intercept of exactly zero when $r_e = 0$, as it must.

| | | | Attenuation slope | | | Intercept | |
|---|---|---|---|---|---|---|---|
| Scenario | $K$ | $r_e$ | simulated | closed form | rel. diff. | simulated | closed form |
| S1 asymmetric power | 300 | 0.00 | 0.6064 | 0.6133 | 1.15% | 0.0047 | 0.0000 |
| S2 matched moderate | 300 | 0.00 | 0.5721 | 0.5787 | 1.16% | 0.0041 | 0.0000 |
| S3 overlap $r_e = 0.05$ | 300 | 0.05 | 0.6062 | 0.6133 | 1.18% | 0.0378 | 0.0339 |
| S4 overlap $r_e = 0.10$ | 300 | 0.10 | 0.6010 | 0.6133 | 2.06% | 0.0709 | 0.0677 |
| S5 small $K$ | 100 | 0.00 | 0.4507 | 0.4555 | 1.06% | 0.0016 | 0.0000 |
| S6 large $K$ | 500 | 0.00 | 0.6880 | 0.6897 | 0.25% | 0.0052 | 0.0000 |

Against 42 held-out configurations not used in its construction, the second-order form of main-text Section 2.3 achieves a mean absolute error of 0.0066 and a maximum of 0.0348, with 88% of cells within 0.01. The leading-order, denominator-only form achieves 0.0492 and 0.1709 on the same cells. The slope varies with $K$ even at fixed heritability, from 0.4555 at $K = 100$ to 0.6897 at $K = 500$, which is the first indication that a single genome-wide correction factor cannot be right.

Table S3: **Interval coverage, width and sign determinacy.** Top: coverage and width before and after the filter, against the real data. Bottom: the fraction of simulated intervals excluding zero as a function of true $\rho$, showing near-symmetry in the sign of $\rho$.

| | Coverage | Mean width | Fraction wider than 1 |
|---|---|---|---|
| Simulated, pre-filter | 0.974 | 1.651 | 0.936 |
| Simulated, post-filter | 0.949 | 1.386 | 0.869 |
| Real data, P4 | — | 1.365 | 0.859 |

| True $\rho$ | Spans zero | Sign determinate |
|---|---|---|
| −0.4 | 0.861 | 0.139 |
| −0.2 | 0.915 | 0.085 |
| 0.0 | 0.937 | 0.063 |
| +0.2 | 0.920 | 0.080 |
| +0.4 | 0.877 | 0.123 |
| +0.6 | 0.779 | 0.221 |

Post-filter coverage is 0.949 against a nominal 0.950, so the filter does not break calibration. Determinacy at $\rho = -0.4$ (0.139) and $\rho = +0.4$ (0.123) is near-identical, so the *count* of sign-determinate loci cannot discriminate between a diametric and a convergent architecture, while their *direction* can. Sample overlap breaks the symmetry, raising determinacy on the positive arm only, which is why the sign test of the main text uses the filtered null rather than a 50/50 binomial.

### S5 Supplementary figure

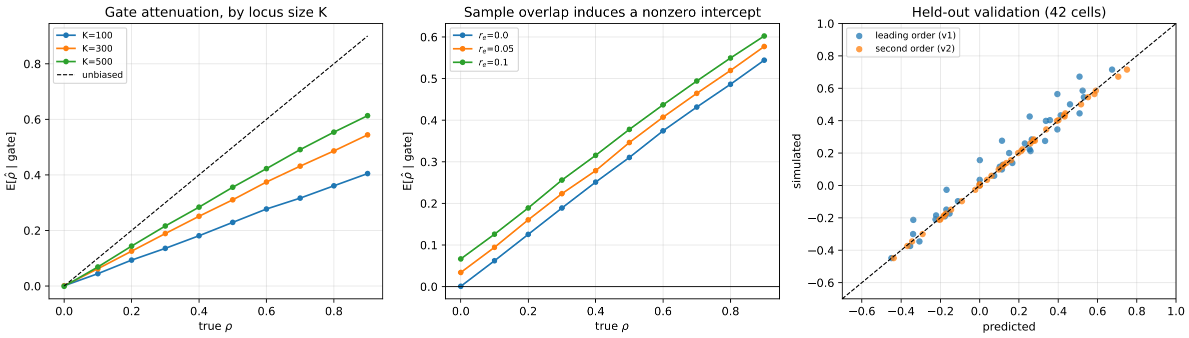


Figure S1: **Tier A diagnostics for the closed-form derivation.** Attenuation of $E[\hat{\rho} \mid G]$ against true $\rho$ at three locus sizes $K$, the same at three levels of sample overlap, and held-out validation of the second-order form against the leading-order one on 42 parameter cells. All panels are sampling-model simulation at 200,000 loci per cell.
